# AAB-seq: An antigen-specific and affinity-readable high-throughput BCR sequencing method

**DOI:** 10.1101/2024.08.13.607736

**Authors:** Mengzhu Hu, Qiuyu Lian, Xiaonan Cui, Xue Dong, Hongyi Xin, Weiyang Shi

## Abstract

B-cell receptor (BCR) sequencing is a powerful antibody discovery tool but current methodology is often inefficient, and lead generation often requires the production and testing of numerous antibody candidates, and it is difficult to provide affinity information for their antibodies at the same time. Here, we introduce AAB-seq (antigen affinity-readable High-throughput BCR sequencing), an efficient antibody screening tool to identify antigen binding affinity of thousands of paired BCRs. It employs fluorophore and DNA barcode-labeled antigen and secondary antibody targeting Ig light chain to label B cells and uses high throughput single cell BCR sequencing and surface protein profiling to obtain the ratio of surface bound antigen to surface BCR in thousands of B-cells. Using AAB-seq, we accurately identified valuable candidate antibodies 1743-3 and 1743-13 from SARS-CoV-2 RBD immunized mouse, providing a basis for further development of SARS-CoV-2 antibody drugs. Thus, AAB-seq allows high throughput identification of antibody sequences paired with antigen affinity, which improves the screening efficiency of functional antibodies and provides an effective solution for the rapid discovery and development of new therapeutic monoclonal antibodies.

## Introduction

Monoclonal antibodies (mAbs) have emerged as a pivotal therapeutic agent for the treatment of oncological and immunological disorders, attributed to their exceptional specificity and capability to potentiate immune responses[1, 2]. Despite advancements in various antibody discovery techniques, the process of identifying and validating functional, high-affinity monoclonal antibodies remain a costly and time-intensive endeavor, typically requiring the screening and assessment of extensive libraries of candidates. Historically, the development of therapeutic antibodies has predominantly relied on methodologies such as hybridoma technology, *in vitro* display techniques, and *ex vivo* single B-cell interrogation, each presenting challenges in terms of either efficiency, affinity, or throughput[3].

Recent advances of single cell sequencing techniques have revolutionized the field of antibody discovery. Each B cell is distinguished by a unique BCR (B cell receptor) gene pair, the sequences of which can be elucidated by sequencing the mRNA from individual B cells. While a few single B cell sequencing technologies have been developed [4-7], they have not been widely adopted in antibody development because BCR sequences, in isolation, lack information on antigen affinity. Consequently, additional procedures need to be developed to recover this critical information, or alternatively, the BCRs sequences need to be validated by traditional biochemical processes or computational prediction. To overcome this, antibody screening platforms that can perform affinity analysis before single B cell isolation and sequencing such as Beacon[8] and Celli*GO* [9]have been developed, but each requires expensive equipment while not being able to perform high-throughput antibody phenotypic analysis.. An alternative approach, LIBRA-seq (Linking B cell Receptor to Antigen Specificity through Sequencing)[10], is able to concurrently obtain antigen binding with BCR sequences via high-throughput single B cell VDJ sequencing (10× Genomics), enabling the screening of thousands of BCR sequences with corresponding antigen binding specificities. Nonetheless, LIBRA-seq has been primarily applied to obtain domain of antigen binding information[11, 12]. During the COVID-19 pandemic, many neutralizing antibodies have been identified directly from blood samples of recovered patients via these single B sequencing methods with success rate between 20-50%[12-16].

Here, we propose an innovative high-throughput BCR sequencing methodology, termed Antigen-affinity BCR sequencing (AAB-seq). This technique is able to obtain antibody affinity toward desired antigens from thousands of BCRs in a high throughput manner, which greatly enhances the efficiency of antibody screening processes. AAB-seq is applicable to purified antigens and holds potential to greatly speed up the development of therapeutic antibodies.

## Results

Antibody affinity is defined as its binding strength to target antigen. To quantify this attribute, we reasoned that one could simultaneously measure the amount of BCR on B cell surface and the number of antigens bound to BCR, and the ration of antigen/BCR can serve as a proximate to such affinity score. To facilitate this measurement, we utilized a cell surface protein sequencing strategy, specifically CITE-seq, to label the antigen and anti-mouse light chain antibody (AAB-Ab) with distinct barcoded DNA oligonucleotides. Additionally, to enhance the selection of BCR-containing cells, we labeled the antigen with fluorophore. These fluorescently labeled antigen-binding B cells were subsequently enriched using fluorescence-activated cell sorting (FACS) and analyzed through VDJ sequencing employing the 10x Genomics VDJ sequencing platform combined with a surface protein profiling kit (Figure 1). This approach enabled the simultaneous capture of individual full BCR sequences along with cell-associated antigen and anti-light chain antibody oligo barcodes from thousands of B cells. In this experimental setup, antibodies with high antigen-binding capacity are expected to have a high AAB-score defined as the function of the ratio of unique molecular identifiers (UMIs) corresponding to the antigen/antibody barcode for each cell, which are experimentally verified by subsequent FACS and surface protein sequencing analysis.

**Figure 1.**
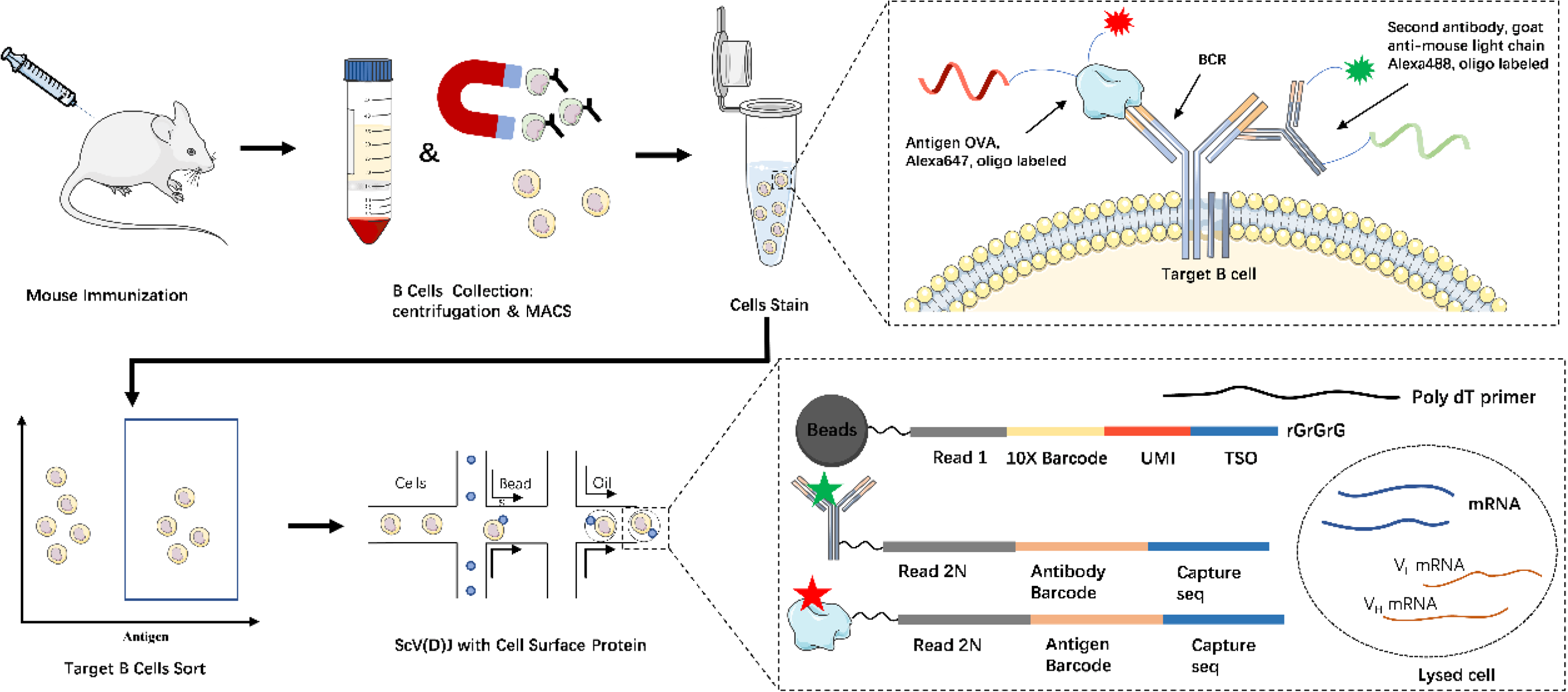
Overview of the workflow for AAB-seq assay. B cells were isolated and enriched from immunized mice by density gradient centrifugation and MACS. And then stained by fluorescently labeled, DNA-barcoded antigen and second antibody, subsequently FACS sorted. Antigen-positive B cells were collected for high-throughput Single Cell V(D)J with Feature Barcode technology. For which, Bead-delivered oligos index both BCR transcripts and antigen/second antibody barcodes during reverse transcription, enabling direct mapping of BCR sequence to antigen/second antibody information following sequencing. The number and placement of oligos and fluorescent molecules on each antigen and second antibody can vary.

### AAB-seq simultaneously obtain antigen-antibody affinity and paired BCR sequence, and AAB-seq helps Rapid and Efficient Screening of Functional Antibodies

To validate the feasibility of AAB-seq, we immunized mouse with chicken ovalbumin (OVA), a well-established model antigen for vaccination studies. We utilized Alexa Fluor 647 and barcoded oligonucleotide-labeled OVA, alongside barcoded AAB-Ab to stain B cells, approximately 10^7 B cells were enriched from the spleens of OVA-immunized mice via density gradient centrifugation and magnetic-activated cell sorting (MACS) targeting CD43, CD4, and Ter-119 markers. To facilitate the identification of B cells, approximately 10^5 B220 PE^+^/DAPI^-^/AF647^+^ cells are sorted and subjected to high-throughput 10x Single Cell V(D)J with Feature Barcode technology according to manufacturer’s suggestions for a target capture of 10,000 B cells (Figure 1, Figure S1, methods).

Following bioinformatic processing, we successfully recovered 3,712 cells with paired heavy-chain and light-chain sequences, as well as antigen reactivity information (Figure 2B, Figure S2A). For each cell, the AAB-score, serving as a critical indicator of antibody affinity, was computed (Methods, Figure 2B). The distribution of AAB-scores ranged from -2.73 to 3.68. To evaluate the correlation between AAB-scores and affinity for each BCR, we selected 16 BCRs for expression and subsequent protein assays (Figures 2A, Figures 2B). We confirmed the predicted antigen specificity and binding capabilities through enzyme-linked immunosorbent assay (ELISA) (Figures S2B), demonstrating that antibodies with high AAB-scores exhibited greater affinity for the antigen, and vice versa (Figures 2A, Figures 2B). Across all tested antibodies, Across all tested antibodies, the correlation between AAB-scores and ELISA-derived area under the curve (AUC) values [(lg(ELISA_AUC)] was robust (Pearson correlation R = 0.8597, P = 9.72E-06) (Figure 2C). These findings indicate that BCR antigen binding affinity can be reliably inferred by measuring the relative ratios of antigen to BCR on B cell surfaces using high-throughput cell surface protein sequencing, thereby providing a direct readout of antigen affinity in single B cell sequencing experiments.

**Figure 2.**
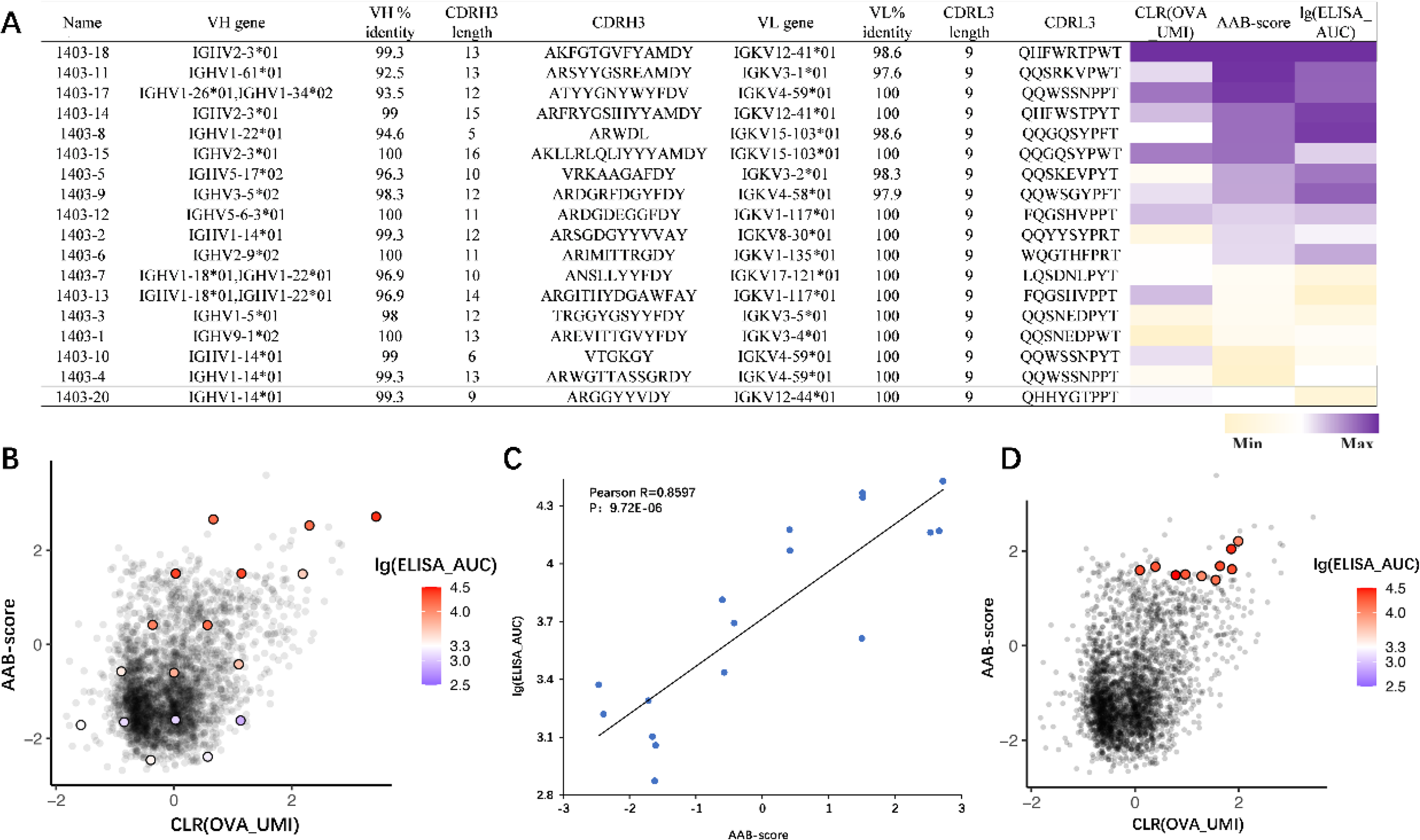
AAB-seq simultaenously obain antigen-antibody affinity and paired BCR sequence and AAB-seq can rapidly and efficiently screen functional antibodies. (A) Sequence characteristics and antigen specificity of dispersedly selected antibodies from OVA immune mice. Percent identity is calculated at the nucleotide level, and CDRH3 and CDRL3 lengths and sequences are noted at the amino acid level. ELISA binding data are displayed as a heatmap of the lg(ELISA_AUC), calculated from data in Figure S2B. Data are represented as mean ± SDs, n≥2. AAB-seq scores and lg(ELISA_AUC) for the selected antibodies are displayed as a heat map from minimum (light yellow) to maximum (purple). (B) All B cells (dots) recovered that had paired heavy/light chain sequencing information and antigen/second antibody reactivity information from the AAB-seq experiment (n = 3,712) are shown, with LIBRA-seq scores for OVA (x axis) and AAB-scores (y axis). Antibodies selected dispersedly for expression and validation (colorized dots) are colored by the LG-AUC scores from lowest (purple) to highest (red). (C) Pearson correlation of AAB-scores (x axis) and ELISA LG-AUC (y axis) for antibodies from OVA AAB-seq experiment; Pearson R= 0.8597, P = 9.72E-06 (two tailed, 95% confidence interval). (D) All B cells (dots) recovered (n = 3,712) are shown as described above. Antibodies with high AAB-score selected (colorized dots) are colored by the LG-AUC scores, the ruler is consistent with (B).

It’s well-known that high antigen binding on B cells can result from either elevated BCR levels or high BCR affinity, thus when antibodies are ranked solely based on the bound antigen amount, as in LIBRA-seq’s scoring system, true antibody binding affinity is not accurately represented. In contrast, the AAB-score provides a precise reflection of antibody affinity for the antigen. Leveraging this insight, we utilized the BCR dataset to identify high-affinity OVA-specific antibodies. B cells with high AAB-scores, as determined from the above high-throughput experiments, were selected for subsequent recombinant antibody expression and characterization (Figures 2D). It is noteworthy that these selected B cells exhibit OVA antigen-positive characterization. We analyzed the sequences of 10 selected antibodies for expression and ested their antigen specificity using ELISA (figure 2D, figure S2D, figure S2C). The results revealed that all antibodies with high AAB-scores bound to OVA and exhibited high antigenic affinity (figure 2D, figure S2C). This validates that functional antibodies can be efficiently screened directly from sequencing data in a targeted manner, thus conserving the resources typically required for expressing and purifying large antibody pools for ELISA tests.

### Discovery of SARS-COV-2 RBD antibodies by AAB-seq

To illustrate the utility of AAB-seq in functional antibody screening, we analyzed the antibody repertoire of mice immunized with the SARS-CoV-2 receptor-binding domain (RBD). As an important immunogenicity region of S protein, RBD contains multiple B-cell and T-cell epitopes and can elicit strong protective antiviral immunity[17]. Following sequencing and subsequent bioinformatic analysis, we recovered paired VH:VL antibody sequences with corresponding AAB-scores for 4,143 cells (figure 3A). From this dataset, we selected 10 antibodies for recombinant antibody production (figure 3A, figure S3A, figure S3B). The purified mAbs were tested for SARS-CoV-2 RBD reactivity by ELISA and biolayer interferometry (BLI), and 8 mAbs with high AAB-scores (1743-3, 1743-6, 1743-8, 1743-9, 1743-10, 1743-11, 1743-12, 1743-13) were identified bind to the RBD and exhibited strong antigen-binding capacities (figure 3A, figure S3C), with BLI results showing their affinity Kd values ranging from 0.211 to 7.80nM (figure S3D, figure S3E), achieving a 100% success rate in antibody screening. Similarly, 2 antibodies with low AAB scores showed no binding to the antigen. Subsequently, we tested these antibodies for their neutralizing activity against the wild-type SARS-CoV-2 pseudovirus. The results indicated that antibodies 1743-3 and 1743-13 effectively neutralized the SARS-CoV-2 WT strain, with IC_50_ values of 2.25μg/mL and 18.58μg/mL, respectively (figure 3B, figure S4A). Next, we assessed the antibody-dependent cellular phagocytosis (ADCP)[18] function of these antibodies against the SARS-CoV-2 RBD protein *in vitro* (methods). Although the results in the pseudovirus neutralization experiments were suboptimal, antibodies 1743-8/9/10/11/12/13 demonstrated good ADCP activity (figure 3C, figure S4B, figure S4C). Finally, antibodies 1743-3 and 1743-13 also exhibited effective antibody-dependent complement deposition (ADCD)[19] (figure 3C, figure S4B, figure S4C). These findings reflect the different profiles of F(ab) and Fc effector functions of the antibodies. These results underscore the effectiveness of AAB-seq in identifying high-affinity antibodies, thereby demonstrating its practical applicability in screening for functional antibodies.

**Figure 3.**
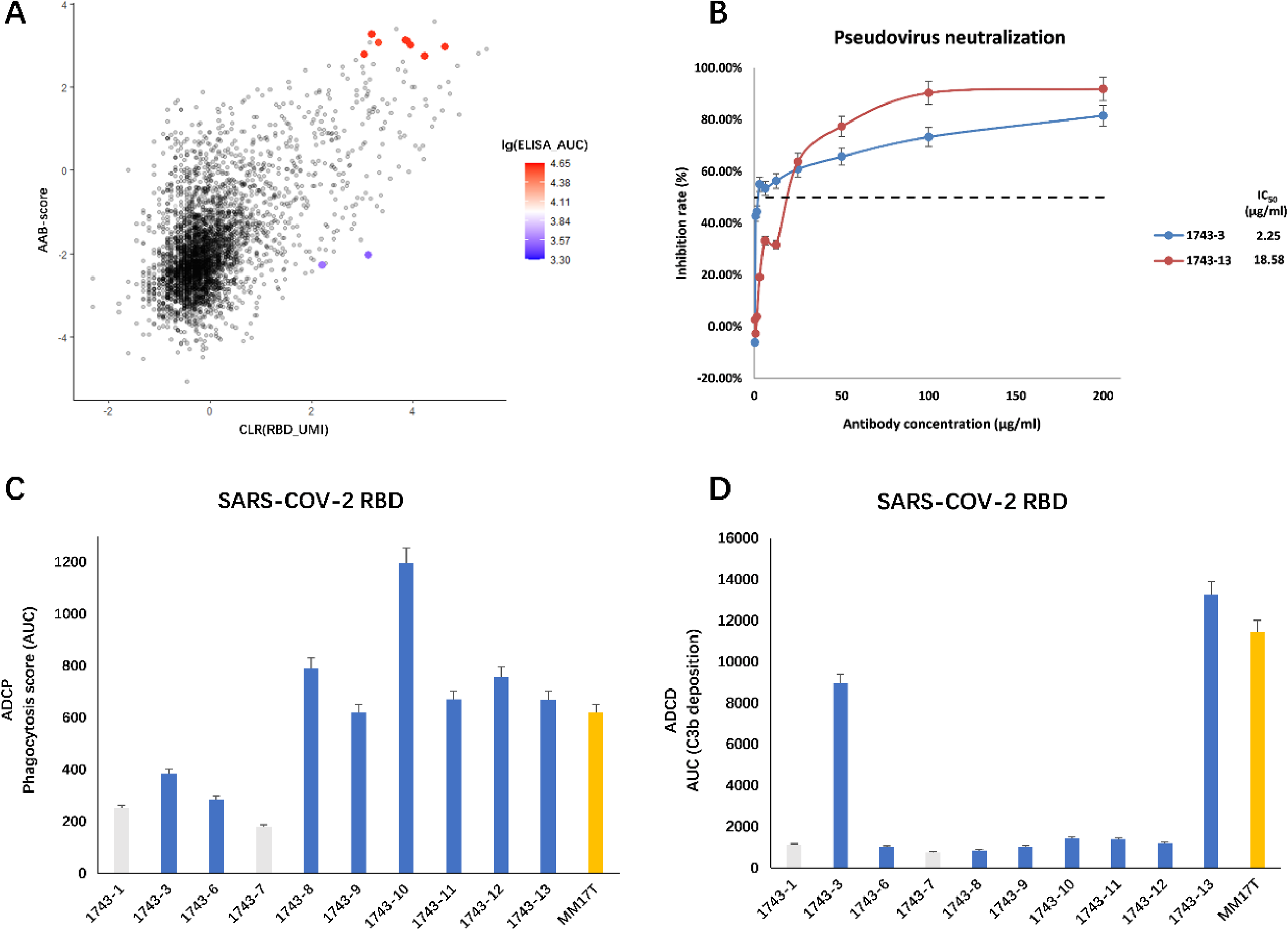
AAB-seq for RBD Antibody Screening and Functional Detection. (A) All B cells (dots) recovered that had paired heavy/light chain sequencing information and antigen/second antibody reactivity information from the AAB-seq experiment for SARS-COV-2 RBD antibodies (n = 3,712) are shown, with CLR(RBD_UMI) (x axis) and AAB-scores (y axis). Antibodies selected dispersedly for expression and validation (colorized dots) are colored by the LG(ELISA_AUC) scores from lowest (purple) to highest (red). (B) Neutralization assays were performed using pseudotyped viruses displaying the S protein from indicated SARS-CoV-2 WT. The values are means ± SD, n = 3. (C) Antibodies were tested for antibody-dependent cellular phagocytosis activity (ADCP) against SARS-CoV-2 RBD, compared to positive control MM17T. AUC of the trogocytosis score is shown, calculated from data in Figure S4B. Data are represented as means ± SDs, n=3. (D) Antibodies were tested for antibody-dependent complement deposition (ADCD) activity against SARS-CoV-2 RBD, compared to positive control MM17T. AUC of the C3b deposition score is shown, calculated from data in Figure S4D. Data are represented as means ± SDs, n=3.

## Discussion

Currently, none of the existing antibody discovery technologies can perfectly achieve high-throughput co-measurement of antibody affinity and BCR sequences. This gap often results in the mass production and expression of many antigen-agnostic B-cell antibodies. Whereas methods capable of analyzing antibody affinity lack high throughput, we have developed a novel high-throughput B-cell screening method AAB-seq that interrogates the antigen affinity of antibodies using sequencing-based readouts. This approach significantly enhances the efficiency of isolating high-affinity antibodies from candidate pools in monoclonal antibody discovery, leading to more precise and effective antibody screening.

To validate our method, we utilized protein OVA as an antigen and demonstrated a strong correlation between AAB-seq scores and ELISA results, and achieving a remarkable screening efficiency of 100%. This represents a significant enhancement compared to the previously methods, which reported efficiencies ranging from 8.8% to 64%[13-15, 20]. In our high-throughput experiments targeting the SARS-CoV-2 RBD, we successfully identified effective candidate RBD antibodies, necessitating only 8 antibodies for subsequent production and validation. This is a substantial improvement over the screening efficiency reported in the literature, which typically necessitates screening hundreds to thousands of candidate antibodies when using RBD as an antigen bait.

We established stringent criteria for the optimal selection of antibodies as follows: (1) Select antigen-positive cells [unique molecular identifier (UMI) > 50]. (2) Select AAB-antibody-positive cells [unique molecular identifier (UMI) > 50]. (3) Remove RBD_UMI abnormal cells (very high UMI data, excludes antigen protein coagulation). (4) Utilize multiple analytical tools to evaluate the antibody sequence, and exclude sequences predicted to be non-productive. (5) Select high AAB-score antibodies for recombinant expression. These specifications ensure the accuracy and efficiency of identifying high-affinity antibodies suitable for further development.

As AAB-seq utilizes DNA barcode oligonucleotides to measure protein binding, it inherently possesses multiplexing capabilities and is not confined to single antigens. Analogous to LIBRA-seq, AAB-seq can theoretically be multiplexed, allowing for the simultaneous screening of antibodies with high binding capacities against multiple antigens. For instance, by concurrently using pneumonia-related coronavirus proteins such as SARS-CoV-S, SARS-CoV-2-S, and MERS-S in experiments, it is possible to obtain both specific antigen-targeting antibodies and broad-spectrum antibodies simultaneously. This innovative approach holds the potential to redefine the standard protocol for single B cell antibody screening.

## Methods

### Conjugation of oligonucleotide barcodes to proteins

We used oligonucleotides that possessed a 15-base pairs protein barcode, a sequence capable of annealing to the template switch oligonucleotide that is part of the 10x bead-delivered oligonucleotides and contains truncated TruSeq small RNA read 1 sequence in the following structure: 5′-CGGAGATGTGTATAAGAGACAG-NNNNNNNNNN-XXXXXXXXXXXXXXX-NNNNNNNNN-CCCATATAAGA*A*A-3′, where Xs represent the protein barcode. The barcodes included OVA (Sigma-Aldrich A5503) (GACTACATAGGCTCC), SARS-COV-2 RBD (2019-nCoV Spike(S) Protein RBD, His-Tag) (CATTGTCGACCGCGA), LC (AffiniPure Goat Anti-Mouse IgG, light chain specific, Jackson ImmunoResearch, Code: 115-005-174) (TAATACCGGAGCCGA, GCACAGTCCGCAATC). Oligos were ordered from Sangon Biotech and IDT with a 5’ amino modification and HPLC purified. In particular, 5′-sulfhydryl-oligonucleotides were conjugated directly to each antigen using Sulfo-SMCC, No-Weigh Format (Thermo Fisher Scientific, A39268) according to manufacturer’s instructions. Briefly, the oligo and protein were desalted, and then protein-NH2 were modified with Sulfo-SMCC 30 minutes at room temperature, and then mixed with oligo-SH 30 minutes at room temperature for the final conjugation after desalted.

### Fluorescent labeling of proteins

After conjugation, the barcoded antigens were mixed with Alexa Fluor™ 647 NHS esters (Thermo Fisher Scientific, A37573) and the barcoded antibodies were mixed with Alexa Fluor™ 488 NHS esters (Thermo Fisher Scientific, A20000), incubate the reaction mixture at room temperature for 2 hours. Add 1M glycine pH 8.5 (final concentration ∼20mM) and incubate at room temperature for 5 minutes to quench residual NHS groups. AKTA FPLC was used to remove excess oligonucleotide from the protein-oligo conjugates. The concentration of the antigen-oligo conjugates was determined by NanoDrop ONE (Thermo Scientific).

### Immunization of mouse and cell sorting

6-8 months old female BALB/c mice (BALB/c) were immunized with 70μg OVA (Sigma-Aldrich, A5503) or SARS-COV2-RBD in Complete Freund’s adjuvant (BD, Franklin Lakes, NJ) followed by biweekly boosts of 35μg OVA or SARS-COV2-RBD in incomplete Freund’s adjuvant divided in three sites (intraperitoneally, subcutaneously at base of tail, at nape of neck and in both hocks). Extraction of cells from spleen 2 days after the last immunization. Single-cell suspensions were prepared by crushing spleen through 40μm nylon mesh sterile cell strainers (Corning, REF352340) and pelleted at 300g for 5min, red blood cells lysed with Lysing Buffer (BD Pharm Lyse™ Lysing Buffer, 555899) according to the manufacturer’s protocol. Then cells washed twice with MACS staining buffer (PBS, 0.5% BSA, and 2mM EDTA) and resuspended in 3mL of MACS buffer, and enriched by negative selection mouse B cell isolation kit (Miltenyi Biotec, #130-090-862, Bergisch Gladbach, Germany) according to the manufacturer’s protocol. Then cells were resuspended in FACS staining buffer at 5 × 10^7^ cells/ml, stained with a cocktail of fluorochrome conjugated anti-mouse B220 PE (BD Biosciences, San Jose, CA) AF488 labeled anti-mouse IgG light chain and AF647 labeled antigen (OVA or SARS-COV-RBD) for sorting of antigen positive B cells on a Beckman Coulter CytoFLEX SRT FACS cell sorter (Supplementary Fig. 1).

### Sample preparation, library preparation and sequencing

After FACS sorting to tube, cells were spun down, resuspended in D-PBS (Sigma-Aldrich) and subjected to cell quality control using TC20 Automated Cell Counter (Bio-rad, 145010). Cell suspensions were loaded onto the Chromium Controller microfluidics device (10x Genomics) and processed using the B cell Single Cell V(D)J solution according to the manufacturer’s suggestions for a target capture of 10,000 B cells, amplify and purify the antigen barcode libraries. Gel Beads-in-Emulsion (GEMs) were formed in channels of a chip and then collected for GEM reverse transcription (GEM-RT) reaction. cDNA was amplified, and additive primers were added to increase the yield of antigen-derived transcript products. After cDNA amplification, the antigen-derived transcript products were size separated from the mRNA-derived cDNA products using SPRI selection and further purification (per the manufacturer’s protocol). After purification, the antigen-derived transcript sequencing library was prepared using a PCR reaction and purified using SPRI purification. The antigen and VDJ libraries were then analyzed, quantified and sequenced using the Illumina NovaSeq platform.

### Sequence processing and bioinformatic analysis

We used Cell Ranger multi (10x Genomics, v6.0.2) to map the sequencing data to the GRCm38 reference genome and generated a matrix of barcode-UMI, barcode-antigen for each sample from paired-end FASTQ files of oligonucleotide libraries as input. Protein barcode libraries were also processed using Cell Ranger (10X Genomics). The BCR data was also processed using cellranger multi with the same version of reference genome. The overlapping barcodes were kept for downstream analysis. R package Seurat (v4.1.1)[21] was used to perform integrated analysis on the cell barcode-UMI-antigen matrix and BCR sequences. Cells were discarded if they met any one of the following criteria: percentage of mitochondrial counts > 25%; number of unique features < 50; number of unique features > 3000. We removed cell barcodes that had only non-functional heavy chain sequences as well as cells with multiple functional heavy chain sequences and/or multiple functional light chain sequences, reasoning that these may be multiplets. Antigen counts for each cell were clr-transformed using compositions package (v2.0-1) [22].

### Determination of AAB-seq Score

Upon obtaining the surface protein UMI count matrix, the noise of UMI<4 is removed. Unique and complete pairs of heavy- and light-chain variable domains are then filtered. Given that UMI serves as a parameter for the actual number of cell-binding proteins, we explicitly define the AAB-score as AAB-score = ln(x1/x2), where x represents the surface protein UMI count, and x1 and x2 correspond to the antigen and AAB-antibody, respectively. While UMI counts for each protein may differ in scale, potentially due to variances in oligonucleotide loading during conjugation, the distribution peaks of a specific protein in the cell population remain consistent under identical conditions. Therefore, the UMI ratio can accurately reflect the relative abundance of two proteins bound to the surface of a single cell. The final AAB-scores are derived by transforming, scaling, and correcting these values using the natural logarithm (LN) function.

### Antibody expression and purification

For each antibody, variable genes were inserted into plasmids encoding the constant region for the heavy chain (pFUSEss-CHIghG1, Invivogen) and light chain (pFUSE2ss-CLIg-hl2, Invivogen and pFUSE2ss-CLIg-hk Invivogen) and synthesized from GenScript. mAbs were expressed in CHO cells. After transfection, cells were cultured at 37°C with 8% CO2 saturation and culture for 5-7 days, and then cell cultures were centrifuged at 6000rpm for 20min. Filtered supernatant was run over a column containing Protein G resin that had been equilibrated with PBS. The column was washed with PBS, and then antibodies were eluted with 0.1M glycine-HCl buffer, pH2.7. After neutralization with 1:10 volume of 1M Tris-HCl pH8, purified antibodies were buffer exchanged into NaAc, 0.2M L-Arginine, pH5.5 using 50kDa Amicon Ultra centrifugal filter units.

### Enzyme linked immunosorbent assay (ELISA)

To assess antibody binding, soluble OVA proteins was plated at 2μg/mL, overnight at 4°C. The next day, plates were washed three times with PBS supplemented with 0.05%Tween-20 (PBS-T) and coated with 5%BSA (Sigma-Aldrich, V900933) in PBS-T. Plates were incubated for 1h at 37°C and washed three times with PBS-T. Primary antibodies were diluted in 0.1%BSA in PBS-T, starting at 10μg/mL with a serial 1:3 dilution and added to the plate. The plates were incubated at 37°C for 1h and washed three times in PBS-T. The secondary antibody, goat anti-mouse IgG-heavy and light chain Antibody conjugated to HRP, was added at a 1:20,000 dilution in 0.1%BSA in PBS-T to the plates, which were incubated for 1h at 37°C. Plates were washed four times with PBS-T and developed by adding TMB substrate (Biolegend, 421101) to each well. The plates were incubated at room temperature for 15min, and 2N sulfuric acid was added to stop the reaction. Plates were read at 450nm. Data are represented as mean ± s.e.m. for one ELISA experiment. ELISAs were repeated two or more times. The AUC was calculated using Origin 2017.

### Biolayer interferometry (BLI) analysis

The BLI-based kinetic assay of antibodies was performed according to a published protocol[23]. Briefly, antibodies to be tested were diluted to the concentration of 5μg/mL with PBS containing 0.02%Tween 20 (Solarbio Life Sciences) and 0.1% (w/v) BSA, and then immobilized onto Anti-mIgG Fc Capture (AMC) biosensors (Sartorius). After a 60-seconds washing step with PBST, biosensor tips were immersed into the wells containing SARS-CoV-2 RBD protein (Sino Biological) diluted to the concentration of 200nM and allowed to associate for 300 seconds, followed by a dissociation step of 600 seconds. The KD were calculated using a 1:1 binding model in Data Analysis Software 11(ForteBio).

### Pseudovirus neutralization assay

The pseudovirus neutralization assay was performed using 293T cells stably expressing ACE2. The cells (30000 cells per well in DMEM with 10% fetal bovine serum) were seeded on a 96-well plate overnight. Various concentrations of mAbs (double serial dilutions starting at 200μg/mL, 50μL aliquots repeats) were mixed with the same volume of SARS-CoV-2 pseudovirus in a 96 well-plate. The mixture was incubated for 1 hour at 37°C with 5% CO2. No-virus control wells were supplied with 100μL of DMEM [10% (v/v) FBS]. Virus-only control wells contained 50μL of medium and 50μL of pseudovirus. After 1 hour, medium was removed from 293T cells, and then, 100μL of the pseudovirus and antibody mixture was incubated with the cells for 1 hour at 37°C with 5% CO2. Another 100μL of DMEM was added into each well and incubated with the cells for 48 hours at 37°C with 5% CO2. After the incubation, supernatants were removed, and 100μL of Nano-Glo Luciferase Assay Reagent (Promega) (1:1 diluted in PBS) was added to each well and incubated for 5 min. Luminescence was measured using a Centro LB 963 Microplate (Berthold Technologies). The relative luciferase unit was calculated by normalizing luminescence signal to the virus-only control group. IC50 was determined by a four-parameter nonlinear regression using GraphPad Prism 9.0 (GraphPad Software Inc.).

### Antibody-dependent cellular phagocytosis (ADCP)

Antibody-dependent cellular phagocytosis (ADCP) was performed using biotinylated SARS-CoV-2 RBD coated fluorescent neutravidin beads as previously described[18]. Briefly, beads were incubated for two hours with antibodies at a starting concentration of 50 mg/mL and titrated fivefold. MM17T (Sino biological, Cat:40592-MM17T) was used as a positive control. Antibodies and beads were incubated with THP-1 cells overnight, fixed and interrogated on the Beckman Coulter CytoFLEX SRT. Phagocytosis score was calculated as the percentage of THP-1 cells that engulfed fluorescent beads multiplied by the geometric mean fluorescence intensity of the population in the FITC channel. For ease of presentation, these scores were then divided by 10000.

### Antibody-dependent complement deposition (ADCD)

Antibody-dependent complement deposition was performed as previously described[19]. Briefly biotinylated SARS-Cov-2 RRBD protein was coated 1:1 onto fluorescent neutravidin beads for 2 hours at 37 degrees. These beads were incubated with 100ug/ml of antibody for 1 hour and incubated with guinea pig complement diluted 1 in 50 with gelatin/veronal buffer for 15 minutes at 37 degrees. Beads were washed at 2000 g twice in PBS and stained with anti-guinea pig C3b-FITC, fixed and interrogated on a Beckman Coulter CytoFLEX SRT. Complement deposition score was calculated as the percentage of C3b-FITC positive beads multiplied by the geometric mean fluorescent intensity of FITC in this population. For ease of presentation, these scores were then divided by 1000.

---

**Figure S1.**
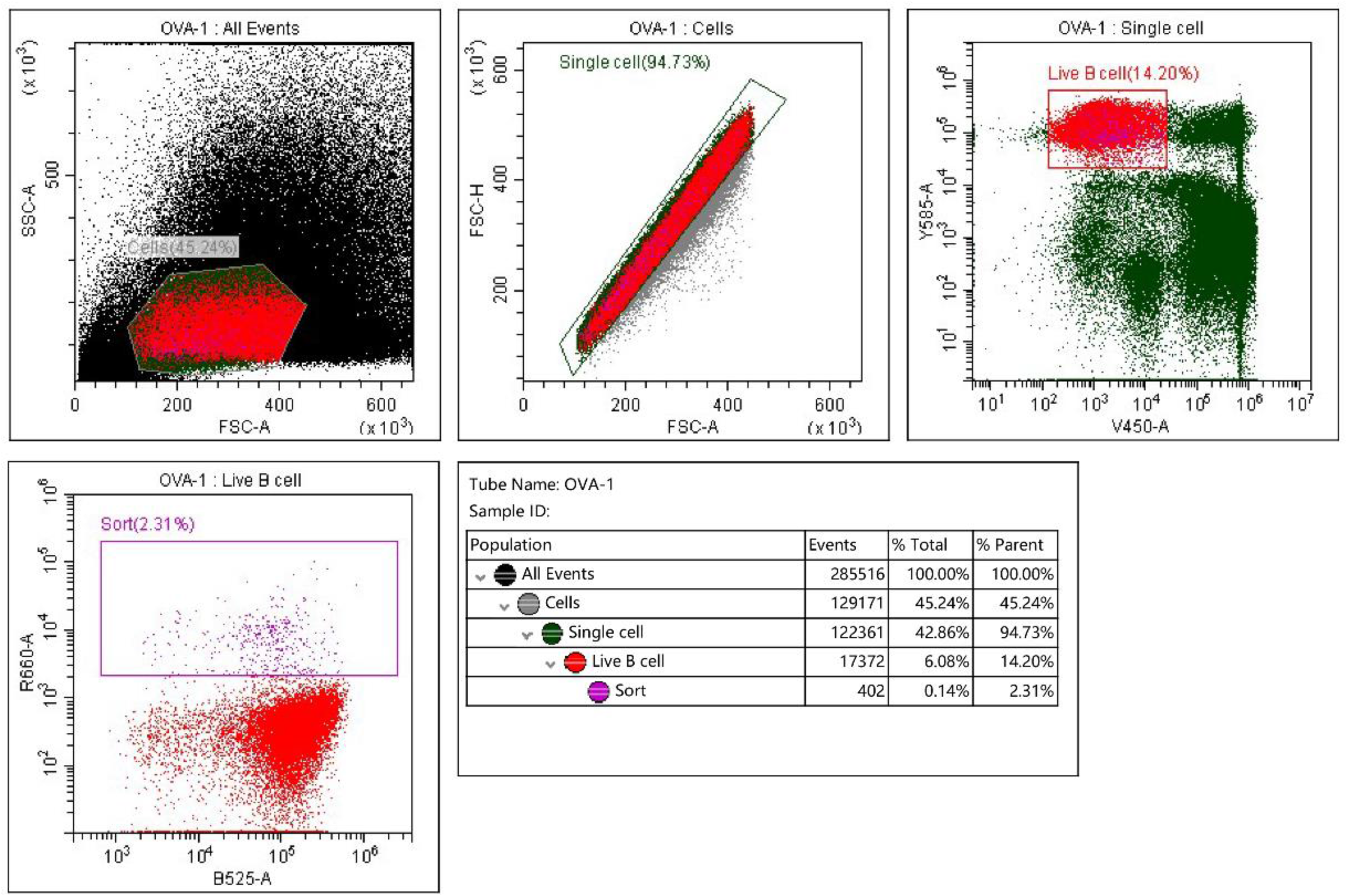
Immunized Balb/c mouse, OVA^pos^ B cell sort gating strategy. 1) FSC vs SSC to gate to exclude fragment, 2) FSC-H vs FSC-A gate to exclude cell doublets, 3) B220-PE vs DAPI for live B cells, 4) Second antibody-AF488 vs OVA-AF647 to sort for OVA^pos^ B cells.

**Figure S2.**
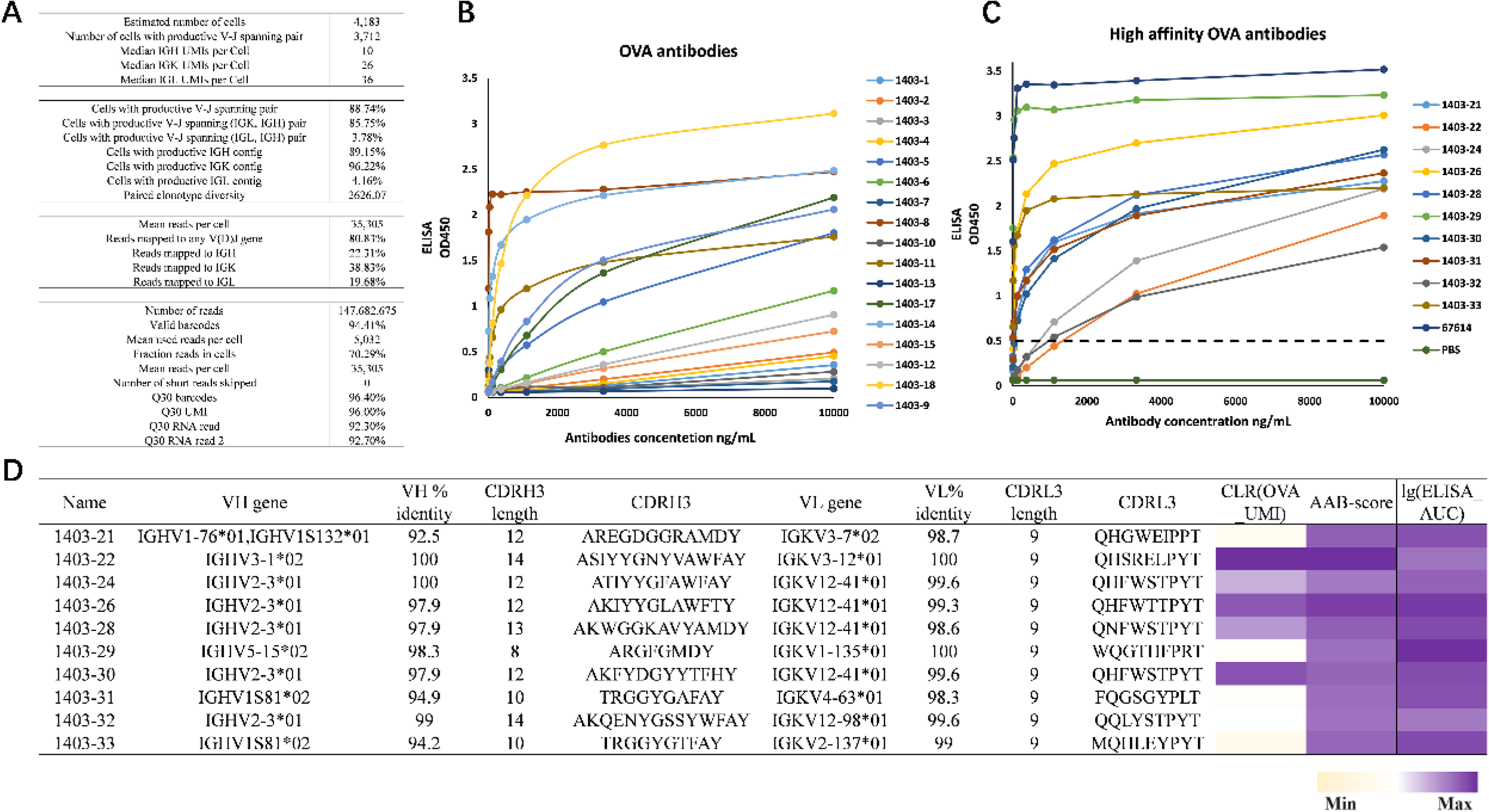
AAB-seq for OVA. (A) The categorization of the number of Cell Ranger-identified (10X Genomics) cells for OVA immune after sequencing is shown. (B) OVA antigen specificity dispersedly antibodies as predicted was validated by ELISA. Data are represented as mean ± SDs, n=3. (C) OVA antigen specificity selected high AAB-score antibodies as predicted was validated by ELISA. Data are represented as mean ± SDs, n=3. (D) Sequence characteristics and antigen specificity of selected high AAB-score antibodies from OVA immune mice. Percent identity is calculated at the nucleotide level, and CDRH3 and CDRL3 lengths and sequences are noted at the amino acid level. ELISA binding data are displayed as a heatmap of the lg(ELISA_AUC), calculated from data in Figure S2C. Data are represented as mean ± SDs, n≥2. AAB-seq scores and lg(ELISA_AUC) for the selected antibodies are displayed as a heat map from minimum (light yellow) to maximum (purple).

**Figure S3.**
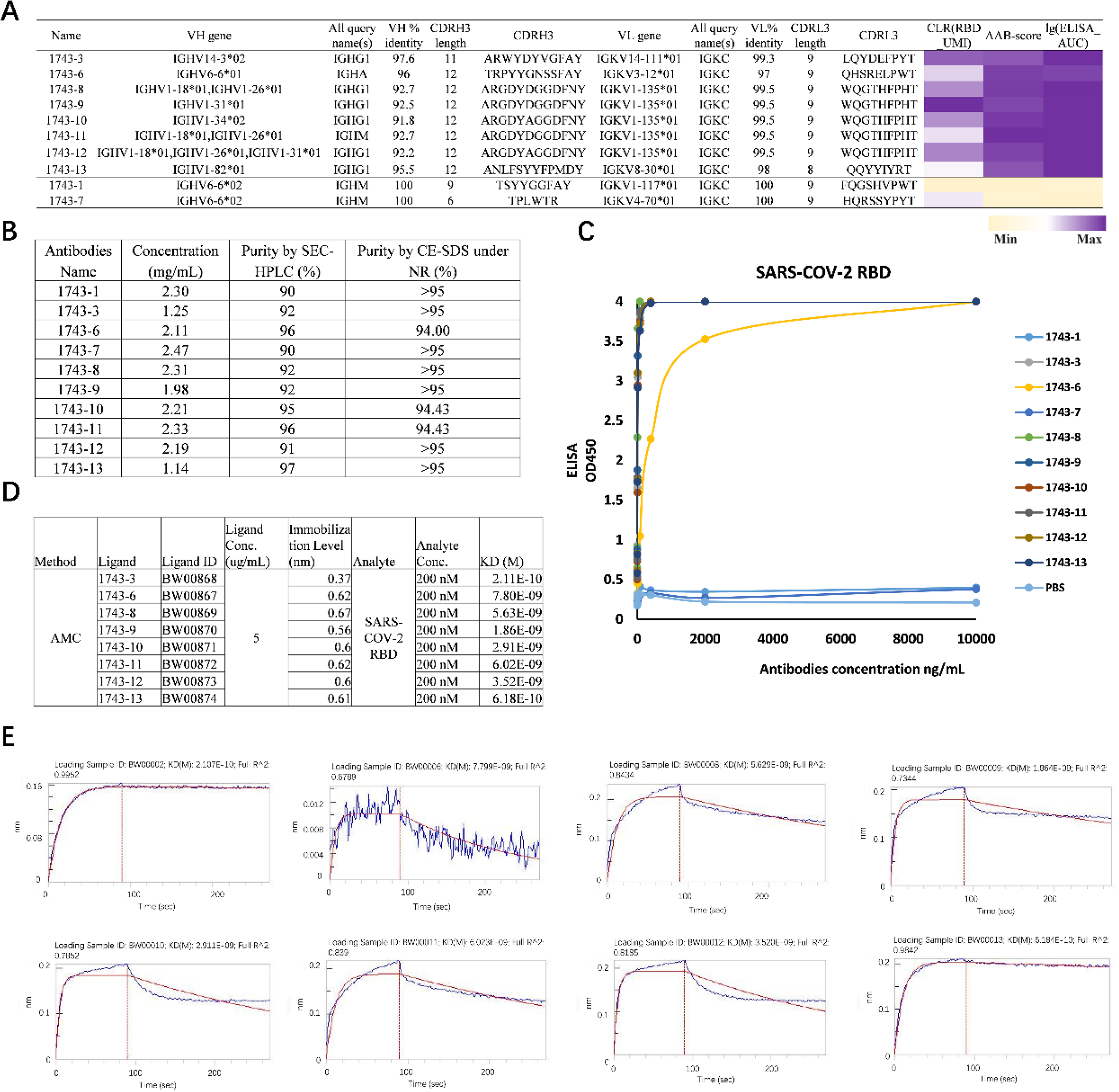
AAB-seq for SARS-COV-2 RBD. (A) Sequence characteristics and antigen specificity of selected antibodies from SARS-COV-2 RBD immune mice. Percent identity is calculated at the nucleotide level, and CDRH3 and CDRL3 lengths and sequences are noted at the amino acid level. ELISA binding data are displayed as a heatmap of the lg(ELISA_AUC), calculated from data in Figure S3C. Data are represented as mean ± SDs, n≥2. AAB-seq scores and lg(ELISA_AUC) for the selected antibodies are displayed as a heat map from minimum (light yellow) to maximum (purple). (B) Recombinant expression, purification, and purity assessment of the selected antibodies. (C) SARS-CoV-2 RBD antigen specificity antibodies as predicted was validated by ELISA. Data are represented as mean ± SDs, n=3. (D) BLI measurement to determine the affinity of the selected antibodies for RBD. (E) The kinetics of binding of selected antibodies to the SARS-COV-2 RBD protein using BLI.

**Figure S4.**
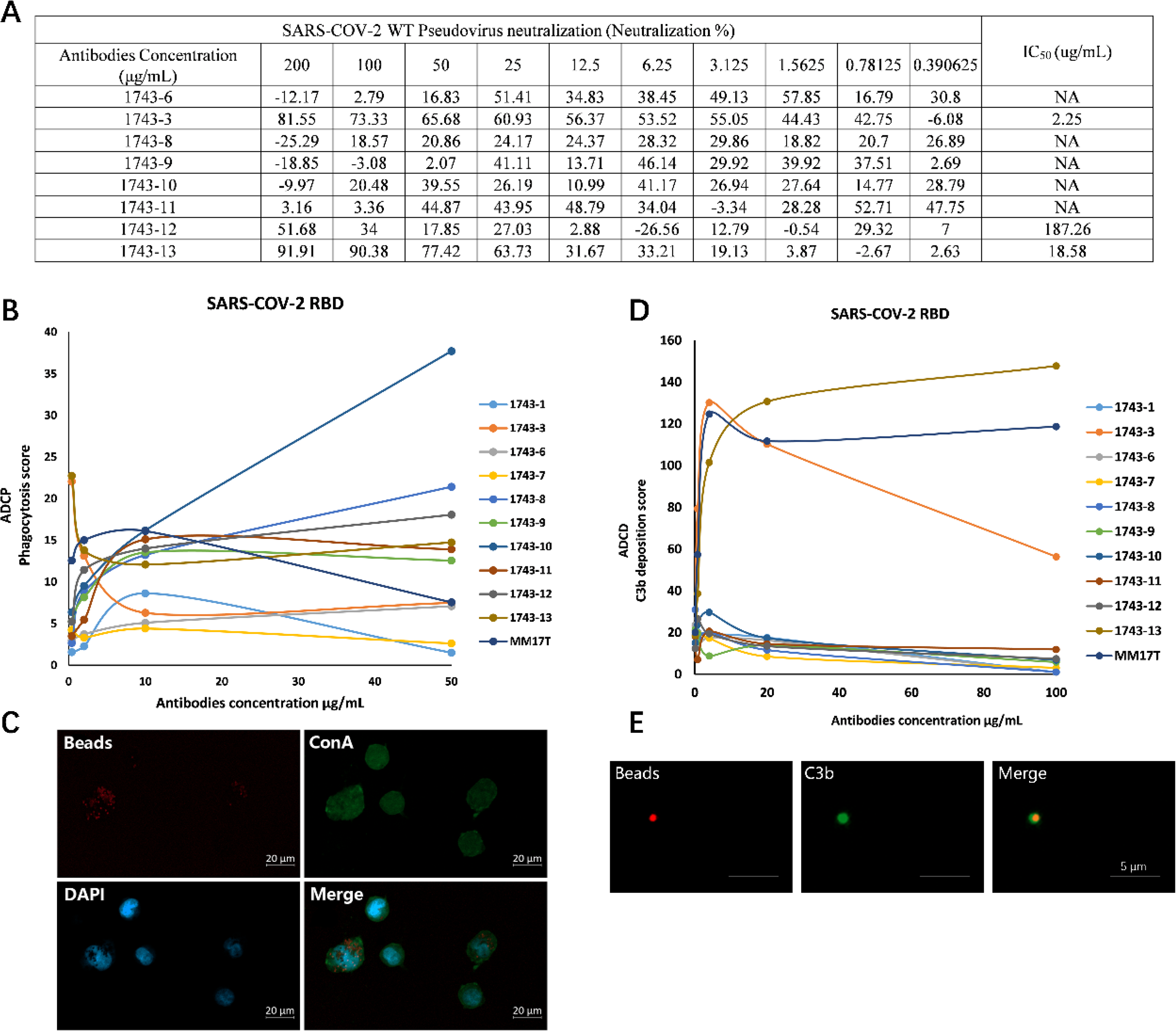
Analysis of SARS-COV-2 RBD Antibody Function. (A) Neutralization assay data of pseudovirus neutralization(%) for SARS-CoV-2 WT. Data are represented as means ± SDs, n=3. (B) Antibodies were tested for antibody-dependent cellular phagocytosis activity against SARS-CoV-2 RBD, compared to positive control MM17T. Phagocytosis score (see Methods) is shown on the y-axis and antibody concentration is shown on the x-axis. Data are represented as means ± SDs, n=3. (C) Confirmation of phagocytosis by Confocal microscopy images. Nuclei are stained blue, beads red, and membrane in green with Concanavalin A(FITC-ConA). (D) Antibodies were tested for antibody-dependent complement deposition (ADCD) activity against SARS-CoV-2 RBD, compared to positive control MM17T. Complement deposition score (see Methods) is shown on the y-axis and antibody concentration is shown on the x axis. Data are represented as means ± SDs, n=3. (E) Confocal microscopy images confirm the detection of complement deposition on fluorescent beads. Beads are stained red, and detection antibody in green with FITC-C3b antibody.

