## supplemental figure for "AAB-seq: An antigen-specific and affinity-readable high-throughput BCR sequencing method"

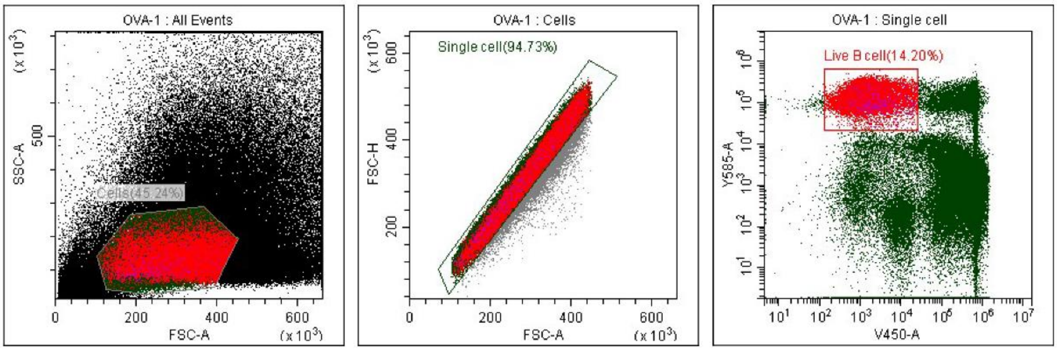


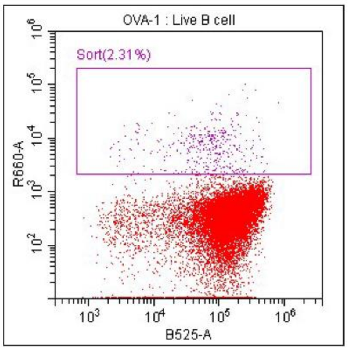


Tube Name: OVA-1
Sample ID:

| Population | Events | % Total | % Parent |
|---|---|---|---|
| All Events | 285516 | 100.00% | 100.00% |
| Cells | 129171 | 45.24% | 45.24% |
| Single cell | 122361 | 42.86% | 94.73% |
| Live B cell | 17372 | 6.08% | 14.20% |
| Sort | 402 | 0.14% | 2.31% |

**Figure S1. Immunized Balb/c mouse, OVA$^{pos}$ B cell sort gating strategy.** 1) FSC vs SSC to gate to exclude fragment, 2) FSC-H vs FSC-A gate to exclude cell doublets, 3) B220-PE vs DAPI for live B cells, 4) Second antibody-AF488 vs OVA-AF647 to sort for OVA$^{pos}$ B cells.

A

| | |
|---|---|
| Estimated number of cells | 4,183 |
| Number of cells with productive V-J spanning pair | 3,712 |
| Median IGH UMIs per Cell | 10 |
| Median IGK UMIs per Cell | 26 |
| Median IGL UMIs per Cell | 36 |
| Cells with productive V-J spanning pair | 88.74% |
| Cells with productive V-J spanning (IGK, IGH) pair | 85.75% |
| Cells with productive V-J spanning (IGL, IGH) pair | 3.78% |
| Cells with productive IGH contig | 89.15% |
| Cells with productive IGK contig | 96.22% |
| Cells with productive IGL contig | 4.16% |
| Paired clonotype diversity | 2626.07 |
| Mean reads per cell | 35,305 |
| Reads mapped to any V(D)J gene | 80.83% |
| Reads mapped to IGH | 22.31% |
| Reads mapped to IGK | 38.83% |
| Reads mapped to IGL | 19.68% |
| Number of reads | 147,682,675 |
| Valid barcodes | 94.41% |
| Mean used reads per cell | 5,032 |
| Fraction reads in cells | 70.29% |
| Mean reads per cell | 35,305 |
| Number of short reads skipped | 0 |
| Q30 barcodes | 96.40% |
| Q30 UMI | 96.00% |
| Q30 RNA read | 92.30% |
| Q30 RNA read 2 | 92.70% |

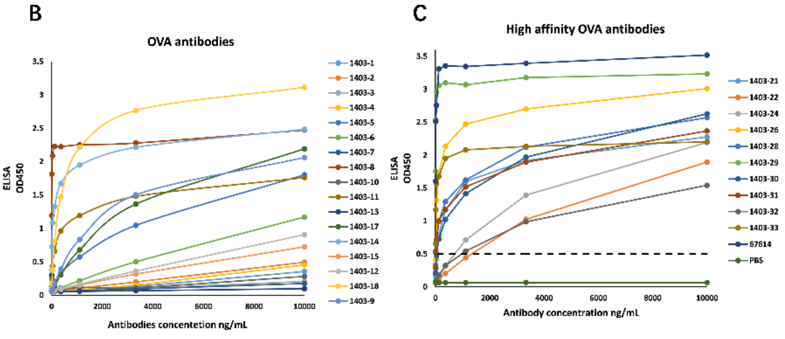


D

| Name | VH gene | VH % identity | CDRH3 length | CDRH3 | VL gene | VL% identity | CDRL3 length | CDRL3 | CLR(OVA_UMI) | AAB-score | lg(ELISA_AUC) |
|---|---|---|---|---|---|---|---|---|---|---|---|
| 1403-21 | IGHV1-76*01,IGHV1S132*01 | 92.5 | 12 | AREGDGGRAMDY | IGKV3-7*02 | 98.7 | 9 | QHGWEIPPT | | | |
| 1403-22 | IGHV3-1*02 | 100 | 14 | ASIYYGNYVAWFAY | IGKV3-12*01 | 100 | 9 | QHSRELPYT | | | |
| 1403-24 | IGHV2-3*01 | 100 | 12 | ATIYYGFAWFAY | IGKV12-41*01 | 99.6 | 9 | QHFWSTPYT | | | |
| 1403-26 | IGHV2-3*01 | 97.9 | 12 | AKIYYGLAWFTY | IGKV12-41*01 | 99.3 | 9 | QHFWTTPYT | | | |
| 1403-28 | IGHV2-3*01 | 97.9 | 13 | AKWGGKAVYAMDY | IGKV12-41*01 | 98.6 | 9 | QNFWSTPYT | | | |
| 1403-29 | IGHV5-15*02 | 98.3 | 8 | ARGFGMDY | IGKV1-135*01 | 100 | 9 | WQGTHFPRT | | | |
| 1403-30 | IGHV2-3*01 | 97.9 | 12 | AKFYDGYYTFHY | IGKV12-41*01 | 99.6 | 9 | QHFWSTPYT | | | |
| 1403-31 | IGHV1S81*02 | 94.9 | 10 | TRGGYGAFAY | IGKV4-63*01 | 98.3 | 9 | FQGSGYPLT | | | |
| 1403-32 | IGHV2-3*01 | 99 | 14 | AKQENYGSSYWFAY | IGKV12-98*01 | 99.6 | 9 | QQLYSTPYT | | | |
| 1403-33 | IGHV1S81*02 | 94.2 | 10 | TRGGYGTFAY | IGKV2-137*01 | 99 | 9 | MQHLEYPYT | | | |

Min Max

**Figure S2. AAB-seq for OVA.**

(A) The categorization of the number of Cell Ranger-identified (10X Genomics) cells for OVA immune after sequencing is shown.

(B) OVA antigen specificity dispersedly antibodies as predicted was validated by ELISA. Data are represented as mean ± SDs, n=3.

(C) OVA antigen specificity selected high AAB-score antibodies as predicted was validated by ELISA. Data are represented as mean ± SDs, n=3.

(D) Sequence characteristics and antigen specificity of selected high AAB-score antibodies from OVA immune mice. Percent identity is calculated at the nucleotide level, and CDRH3 and CDRL3 lengths and sequences are noted at the amino acid level. ELISA binding data are displayed as a heatmap of the lg(ELISA_AUC), calculated from data in Figure S2C. Data are represented as mean ± SDs, n⩾2. AAB-seq scores and lg(ELISA_AUC) for the selected antibodies are displayed as a heat map from minimum (light yellow) to maximum (purple).

A

| Name | VH gene | All query name(s) | VH % identity | CDRH3 length | CDRH3 | VL gene | All query name(s) | VL% identity | CDRL3 length | CDRL3 | CLR(RBD_UMI) | AAB-score | lg(ELISA_AUC) |
|---|---|---|---|---|---|---|---|---|---|---|---|---|---|
| 1743-3 | IGHV14-3*02 | IGHG1 | 97.6 | 11 | ARWYDYVGFAY | IGKV14-111*01 | IGKC | 99.3 | 9 | LQYDEFPYT | | | |
| 1743-6 | IGHV6-6*01 | IGHA | 96 | 12 | TRPYYGNSSFAY | IGKV3-12*01 | IGKC | 97 | 9 | QHSRELPWT | | | |
| 1743-8 | IGHV1-18*01,IGHV1-26*01 | IGHG1 | 92.7 | 12 | ARGDYDGGDFNY | IGKV1-135*01 | IGKC | 99.5 | 9 | WQGTHFPHT | | | |
| 1743-9 | IGHV1-31*01 | IGHG1 | 92.5 | 12 | ARGDYDGGDFNY | IGKV1-135*01 | IGKC | 99.5 | 9 | WQGTHFPHT | | | |
| 1743-10 | IGHV1-34*02 | IGHG1 | 91.8 | 12 | ARGDYAGGDFNY | IGKV1-135*01 | IGKC | 99.5 | 9 | WQGTHFPHT | | | |
| 1743-11 | IGHV1-18*01,IGHV1-26*01 | IGHM | 92.7 | 12 | ARGDYDGGDFNY | IGKV1-135*01 | IGKC | 99.5 | 9 | WQGTHFPHT | | | |
| 1743-12 | IGHV1-18*01,IGHV1-26*01,IGHV1-31*01 | IGHG1 | 92.2 | 12 | ARGDYAGGDFNY | IGKV1-135*01 | IGKC | 99.5 | 9 | WQGTHFPHT | | | |
| 1743-13 | IGHV1-82*01 | IGHG1 | 95.5 | 12 | ANLFSYYFPMDY | IGKV8-30*01 | IGKC | 98 | 8 | QQYYIYRT | | | |
| 1743-1 | IGHV6-6*02 | IGHM | 100 | 9 | TSYYGGFAY | IGKV1-117*01 | IGKC | 100 | 9 | FQGSHVPWT | | | |
| 1743-7 | IGHV6-6*02 | IGHM | 100 | 6 | TPLWTR | IGKV4-70*01 | IGKC | 100 | 9 | HQRSSYPYT | | | |

Min Max

B

| Antibodies Name | Concentration (mg/mL) | Purity by SEC-HPLC (%) | Purity by CE-SDS under NR (%) |
|---|---|---|---|
| 1743-1 | 2.30 | 90 | >95 |
| 1743-3 | 1.25 | 92 | >95 |
| 1743-6 | 2.11 | 96 | 94.00 |
| 1743-7 | 2.47 | 90 | >95 |
| 1743-8 | 2.31 | 92 | >95 |
| 1743-9 | 1.98 | 92 | >95 |
| 1743-10 | 2.21 | 95 | 94.43 |
| 1743-11 | 2.33 | 96 | 94.43 |
| 1743-12 | 2.19 | 91 | >95 |
| 1743-13 | 1.14 | 97 | >95 |

C

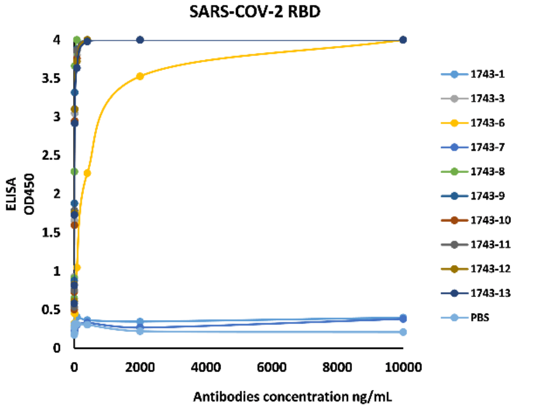


D

| Method | Ligand | Ligand ID | Ligand Conc. (ug/mL) | Immobilization Level (nm) | Analyte | Analyte Conc. | KD (M) |
|---|---|---|---|---|---|---|---|
| AMC | 1743-3 | BW00868 | 5 | 0.37 | SARS-COV-2 RBD | 200 nM | 2.11E-10 |
| | 1743-6 | BW00867 | | 0.62 | | 200 nM | 7.80E-09 |
| | 1743-8 | BW00869 | | 0.67 | | 200 nM | 5.63E-09 |
| | 1743-9 | BW00870 | | 0.56 | | 200 nM | 1.86E-09 |
| | 1743-10 | BW00871 | | 0.6 | | 200 nM | 2.91E-09 |
| | 1743-11 | BW00872 | | 0.62 | | 200 nM | 6.02E-09 |
| | 1743-12 | BW00873 | | 0.6 | | 200 nM | 3.52E-09 |
| | 1743-13 | BW00874 | | 0.61 | | 200 nM | 6.18E-10 |

E

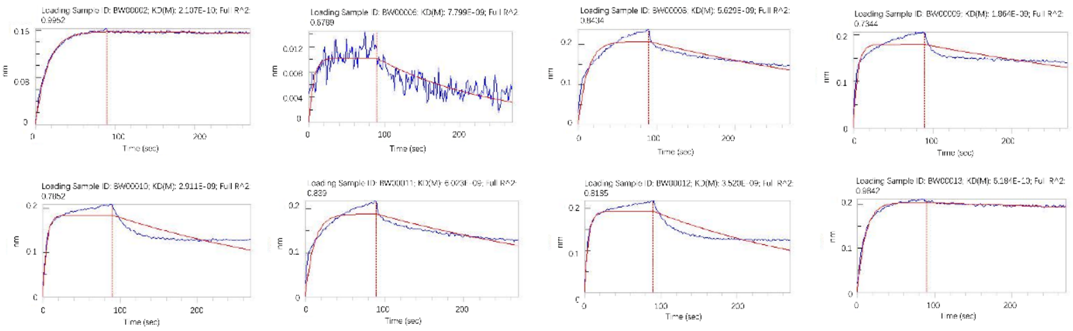


Figure S3. **AAB-seq for SARS-COV-2 RBD.**

(A) Sequence characteristics and antigen specificity of selected antibodies from SARS-COV-2 RBD immune mice. Percent identity is calculated at the nucleotide level, and CDRH3 and CDRL3 lengths and sequences are noted at the amino acid level. ELISA binding data are displayed as a heatmap of the lg(ELISA_AUC), calculated from data in Figure S3C. Data are represented as mean ± SDs, n⩾2. AAB-seq scores and lg(ELISA_AUC) for the selected antibodies are displayed as a heat map from minimum (light yellow) to maximum (purple).

(B) Recombinant expression, purification, and purity assessment of the selected antibodies.

(C) SARS-CoV-2 RBD antigen specificity antibodies as predicted was validated by ELISA. Data are represented as mean ± SDs, n=3.

(D) BLI measurement to determine the affinity of the selected antibodies for RBD.

(E) The kinetics of binding of selected antibodies to the SARS-COV-2 RBD protein using BLI.

A

| SARS-COV-2 WT Pseudovirus neutralization (Neutralization %) | | | | | | | | | | | $IC_{50}$ (ug/mL) |
|---|---|---|---|---|---|---|---|---|---|---|---|
| Antibodies Concentration (μg/mL) | 200 | 100 | 50 | 25 | 12.5 | 6.25 | 3.125 | 1.5625 | 0.78125 | 0.390625 | |
| 1743-6 | -12.17 | 2.79 | 16.83 | 51.41 | 34.83 | 38.45 | 49.13 | 57.85 | 16.79 | 30.8 | NA |
| 1743-3 | 81.55 | 73.33 | 65.68 | 60.93 | 56.37 | 53.52 | 55.05 | 44.43 | 42.75 | -6.08 | 2.25 |
| 1743-8 | -25.29 | 18.57 | 20.86 | 24.17 | 24.37 | 28.32 | 29.86 | 18.82 | 20.7 | 26.89 | NA |
| 1743-9 | -18.85 | -3.08 | 2.07 | 41.11 | 13.71 | 46.14 | 29.92 | 39.92 | 37.51 | 2.69 | NA |
| 1743-10 | -9.97 | 20.48 | 39.55 | 26.19 | 10.99 | 41.17 | 26.94 | 27.64 | 14.77 | 28.79 | NA |
| 1743-11 | 3.16 | 3.36 | 44.87 | 43.95 | 48.79 | 34.04 | -3.34 | 28.28 | 52.71 | 47.75 | NA |
| 1743-12 | 51.68 | 34 | 17.85 | 27.03 | 2.88 | -26.56 | 12.79 | -0.54 | 29.32 | 7 | 187.26 |
| 1743-13 | 91.91 | 90.38 | 77.42 | 63.73 | 31.67 | 33.21 | 19.13 | 3.87 | -2.67 | 2.63 | 18.58 |

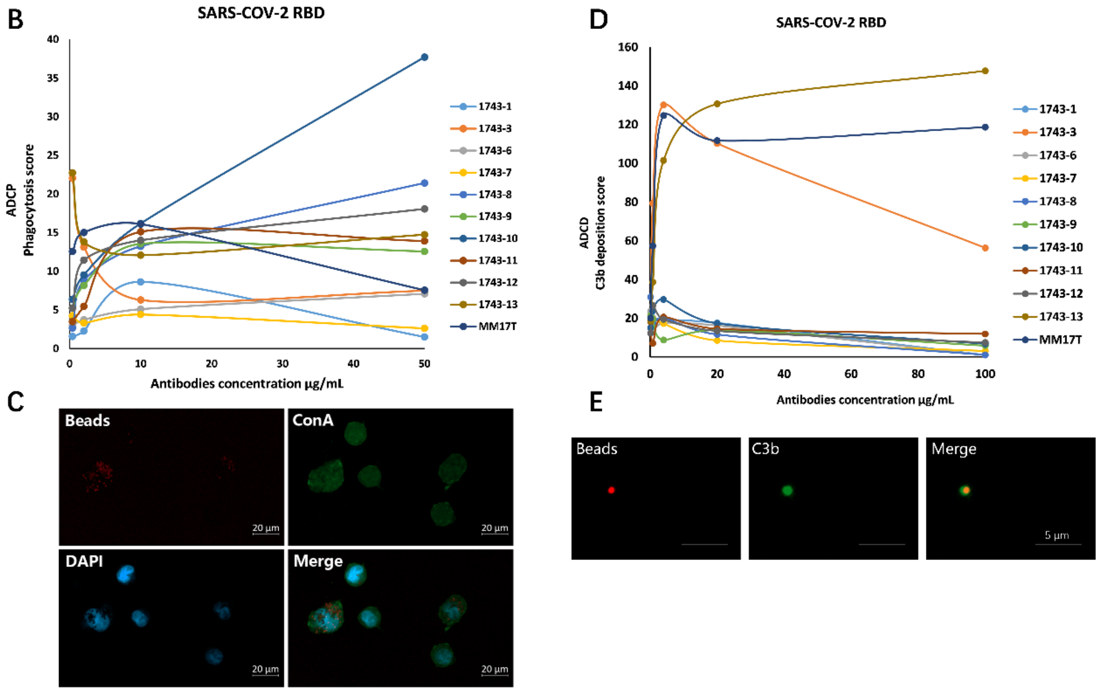


**Figure S4. Analysis of SARS-COV-2 RBD Antibody Function.**

(A) Neutralization assay data of pseudovirus neutralization(%) for SARS-CoV-2 WT. Data are represented as means ± SDs , n=3.

(B) Antibodies were tested for antibody-dependent cellular phagocytosis activity against SARS-CoV-2 RBD, compared to positive control MM17T. Phagocytosis score (see Methods) is shown on the y-axis and antibody concentration is shown on the x-axis. Data are represented as means ± SDs, n=3.

(C) Confirmation of phagocytosis by Confocal microscopy images. Nuclei are stained blue, beads red, and membrane in green with Concanavalin A(FITC-ConA).

(D) Antibodies were tested for antibody-dependent complement deposition (ADCD) activity against SARS-CoV-2 RBD, compared to positive control MM17T. Complement deposition score (see Methods) is shown on the y-axis and antibody concentration is shown on the x axis. Data are represented as means ± SDs , n=3.

(E) Confocal microscopy images confirm the detection of complement deposition on fluorescent beads. Beads are stained red, and detection antibody in green with FITC-C3b antibody.
